# Biological invasions forming intraguild predation communities in homogeneous and heterogeneous landscapes

**DOI:** 10.1101/2024.08.15.608146

**Authors:** Silas Poloni, Roberto André Kraenkel, Renato Mendes Coutinho

**Affiliations:** Institute of Theoretical Physics, São Paulo State University, R. Dr. Bento Teobaldo Ferraz, 271, São Paulo, 01140-070, São Paulo, Brazil; Department of Biology, University of Victoria, 3800, Finnerty Road, Victoria, V8P 5C2, British Columbia, Canada; Center for Mathematics, Computing and Cognition, Federal University of ABC, Av. dos Estados, 5001, Santo André,, 09210-580, São Paulo, Brazil

**Keywords:** Spatial ecology, population dynamics, reaction-diffusion equations, mathematical models

## Abstract

Intraguild predation (IGP) allows for coexistence between two consumers of a single resource, as long as the intraguild prey (IG prey) is competitively superior to the Intraguild Predator (IG predator) and resource population productivity is neither abundant or limiting. Here we explore biological invasions forming IGP community modules by either introducing IG prey or IG predator species to established Consumer-Resource populations in homogeneous and heterogeneous landscapes, using reaction-diffusion equations as our modeling framework. Our main methods of analysis are comparing numerical solutions to linearization techniques and homogenization approximations. We find that in homogeneous landscapes, speeds are linearly determinate, i.e., depend on low invader population densities at the leading edge. We also find traveling wave solutions and dynamical stabilization regimes. On heterogeneous landscapes, our results show that depending on habitat preferences of the three species involved, coexistence regimes can occur regardless of IG-Prey being least effective consumer, or be hindered even when IG-Prey remais as the dominant competitor. Our work asses how fast can organisms invade novel landscapes in presence of a established IG prey or IG predator, and also demonstrates how habitat fragmentation and species habitat preference can disrupt or facilitate coexistence in IGP communities.

## 1 Introduction

Often, the introduction of new consumer species in a novel habitat can generate intraguild predation (IGP) interactions, a multi-species trophic network where exploitative competitors for a shared resource present a predator-prey relation (Polis et al., 1989; Polis and Holt, 1992), and may lead to exclusion of native species, failure of invasion, or the formation of an IGP community (Tuckett et al., 2021; Montserrat et al., 2012; Fritts and Rodda, 1998; Grosholz and Ruiz, 1995). Alongside interspecific interactions, landscape heterogeneity plays an important role on species spatial distributions, coexistence regimes and movement behavior (Polis et al., 1997; Schtickzelle and Baguette, 2003; Abrams, 2007), which in turn may also significantly change how invasions occur. Because so many factors can alter the course of range expansion events, spatially structured mathematical models have been vastly used to explore possible outcomes of biological invasions, and significant advances in the field allow us to estimate spreading speeds (see, for instance, Castillo-Chavez et al. (2013)) and analyze heterogeneous landscapes in a simplified manner (see Yurk and Cobbold (2018); Cobbold et al. (2022)). Nonetheless, theoretical approaches for three species IGP communities haven’t been studied in detail in this context, and can reveal regime shifts caused by the invasion of novel consumers, as well as the main factors behind it, such as the interplay of demographic traits and dispersal behaviors. In this work, we study a three species IGP model in different landscape settings and provide spreading speeds estimates, show some of the regimes found numerically, and provide some key factors that change community formation processes.

The formation of IGP relations between local and invader species are not rare. One of the best known invader species to partake in IGP relations with local community species is the harlequin ladybird (*Harmonia axyridis*), either as the IG predator, prey or both in cases of symmetrical IGP (Berg et al., 2012; Ware and Majerus, 2008) (see also Alhmedi et al. (2010) for a review of the topic). Another common IGP formation between local and invasive species is between guppies (*Poecilia reticulata*) and mosquitofish (*Gambusia spp*.). Both guppies and mosquitofish are widely used for mosquito population control, and have been introduced in many places around the globe, often in each others territories (native or previously invaded), their diets constitute in algae, insect larvae and the other species eggs whenever present, yielding a symmetrical IGP (Tuckett et al., 2021; Vera et al., 2016; Tsurui-Sato et al., 2019). These are some examples on introduced species that interact with local communities through intraguild predation, revealing how common this interactions are in nature.

Theoretical approaches to account for population spread have been continuously developed to address different biological and ecological phenomena, mainly because the large temporal and spatial scales usually prevent experiments. The mathematical description of biological invasions, and other spatial ecology phenomena, follows the pioneering works of Fisher (1937); Kolmogorov A et al. (1937) and Skellam (1951), that were the first of many that helped establish reaction diffusion equations (RDE) as the main workhorses of spatial ecology. In this modeling framework, individuals of a given species are assumed to move with Brownian motion (hence diffusion), and population grows/decays according to the relevant biological processes governing demography, such as reproduction, death and intra-specific competition (hence reaction). With this theoretical approach, we have been able to estimate, for example, spreading speeds of invasive species (Hadeler and Rothe, 1975; Weinberger, 1978), and determine the existence of traveling wave solutions, i.e., spatial profiles that are maintained through time, but advance in space, signaling the range expansion and establishment of invading species (Weinberger, 1982).

The mathematical foundations for IGP are given in (Holt and Polis, 1997), where a three species IGP network is considered. There, we have two consumers of a single biotic resource, with a predation relation among themselves, as displayed in figure 1. Such predation relation allows coexistence between both consumers, in cases which otherwise would be unattainable due to competitive exclusions (Tilman and others, 1990; Klausmeier and Tilman, 2002). The precise conditions in which coexistence is possible depend on how large is resource productivity/carrying capacity, and on the intraguild prey (IG prey) being a stronger exploitative competitor than intraguild predator (IG predator) (Holt and Polis, 1997). Following Tilman and others (1990), the later translates into IG prey leading the resource to lower populational levels than IG predator (when each consumer is set with resource alone). Beyond this results, (Holt and Polis, 1997) highlights many venues in which the theory can be pushed forward to explain other potential coexistence mechanisms, such as considering age structure, adaptive behaviors and spatial dynamics, such as dispersal and habitat heterogeneity.

Accounting for inter-specific interactions in RDE frameworks has been the target of many studies, specially in competition and consumer-resource problems (Owen and Lewis, 2001; Petrovskii and Malchow, 2000; Malchow, 1997; Lewis et al., 2002; Hosono, 1998). Since IGP consists of competition and predation, results from such models can provide some expectations for IGP as well. One characteristic RDE models for competition and predator-prey interactions display are the linear determinacy of spreading speeds (see Petrovskii and Malchow (2000); Malchow and Petrovskii (2002); Weinberger et al. (2002); Lewis et al. (2002); Hosono (1998) for the required conditions for linear determinacy). This means that the speed at which an invasive species spreads through the landscape depends solely on low invader density dynamics, where the alien species is still establishing, and resident community is still very close to its equilibrium. Another common finding in this two species systems is the formation of traveling wave solutions ^1^, which, for example, show the displacement of the resident single species state to a coexistence one, as the invader species spreads. Noteworthy, predator-prey dynamics can exhibit various spatial profiles other than traveling wave solutions, such as pattern formation, dynamical stability and chaos, depending on the stability of the coexistence state (in the non-spatially structured model) (Malchow and Petrovskii, 2002; Malchow, 1997; Petrovskii and Malchow, 2001; Murray, 2003).

**Fig. 1:**
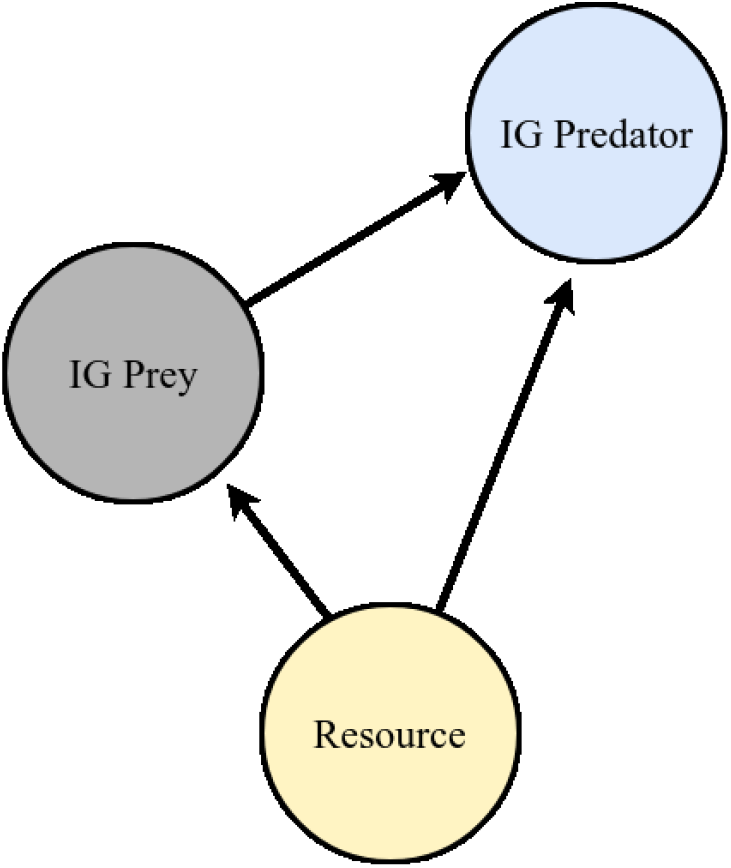
A three species intraguild predation network diagram

While RDE formulations for competition and consumer-resource are well described, spatially structured models for IGP are few, mainly because accounting for more than two species can often generate complex dynamics in space, and provide less informative results. With that, Bampfylde and Lewis (2007) formulate a two species Lotka-Voltera competition model (with added terms for predation) to understand the dynamics of Intraguild Predation. They show that invasions leading to coexistence, extinction and bi-stable regimes are possible, which imply an invasive species may advance in space and exclude the resident species, coexist with it or, sometimes, regress and fail to invade the landscape. In Hall (2010) two different models, one assuming interference competition between IG prey and IG predator (leading to a two species model), and another considering exploitative competition (leading to a three species model), show that successful IG-predator invasions can be even faster than they would in the absence of the IG prey. This happens when the fitness benefits of IG predation outweigh the fitness loss through competition.

Reaction diffusion equations have been also developed and applied to study heterogeneous landscapes in Shigesada et al. (1986), with the landscape being composed of two patch types that are arranged periodically over the real line. Movement from one patch type to the other is such that flux and population densities are continuous, which do not address specifically possible patch preferences of the invasive and/or resident species that inhabit such landscape. To account for such preferences, Maciel and Lutscher (2013) considers individual movement at the interfaces between different patches as being biased towards a preferred patch type, following Ovaskainen and Cornell (2003). This results in interface conditions that are discontinuous in density, but continuous in flux, with population densities potentially accumulating at edges of the preferred patch type.

A method to approximate the results of RDE models in periodic landscapes is presented in Yurk and Cobbold (2018). The main idea is to assume the dynamics inside a pair of patches occur at a much smaller time scale than that of the whole landscape, allowing for the small scale processes to be averaged in the large scale. With this method, Maciel and Lutscher (2018) investigated how movement behaviors can cause regime shifts in competing species, finding that reversals ^2^ can occur whenever the habitat preference of the weakest competitor allows for it to spend an optimal time in each of the habitats, increasing its effective carrying capacity while decreasing the negative effects from competition with the stronger competitor.

Through a multi-patches model, Amarasekare (2007) studies the IGP in discrete space, and provides some insights for the problem of dispersal strategies and habitat selection. In her work, IG prey and predator move between three connected patches of varying resource productivity (resource is stationary). The investigations and results focus on the roles of IG prey and predator dispersal, showing that different strategies lead to different spatial distributions, e.g., fitness based dispersal leads to segregated coexistence, where IG prey stays on low resource productivity patches, and IG predator on high resource productivity ones.

Although some insightful results for IGP in space are present in literature, accounting for both resource population levels and dispersal is lacking, both potential key processes to understanding coexistence and exclusion regimes. Also, measuring speeds of invasion of a consumer species can unravel how fast possible regime shifts take place. Beyond that, we can verify the formation of spatial profiles such as traveling wave solutions and possible oscillatory regimes. Understanding the homogeneous landscape problem can also provide expectations for the large spatio-temporal scale for IGP in heterogeneous landscapes, in which patch preference behavior of resource and consumers populations can be accounted for explicitly, and possibly mediate competitive reversals, which in IGP communities would mean shifting coexistence regimes into exclusion ones and vice-versa (Holt and Polis, 1997; Polis and Holt, 1992).

In this work, we consider a version of the classical IGP model by Holt and Polis (1997) with added ecological diffusion terms to account for movement in homogeneous and heterogeneous landscapes. In section 2, we present the model in homogeneous landscapes, and measure spreading speeds as well as display some of the regimes found. In section 3, we present the corresponding model in heterogeneous/periodic landscapes, following (Yurk and Cobbold, 2018; Cobbold et al., 2022), we perform the homogenization technique, and drawing correspondence with our findings in the homogeneous landscape model, we determine conditions for mutual invasibility in the large spatio-temporal scales. Finally, in section 4, we discuss our results and present future venues of research.

## 2 Intraguild Predation in Homogeneous Landscapes

### 2.1 Model

We consider that population densities vary in continuous time, *t*, and space, *x* and denote IG prey density as *C*_1_ ≡ *C*_1_(*t, x*), IG predator as *C*_2_ ≡ *C*_2_(*t, x*), and the shared resource as *R* ≡ *R*(*t, x*). In our model, every species move with “ecological” diffusion as in Ovaskainen and Cornell (2003); Maciel and Lutscher (2013), predation relations are linear, while consumers are subject to natural mortality and resource grows and reproduces according a density dependent growth function. The model equations are then

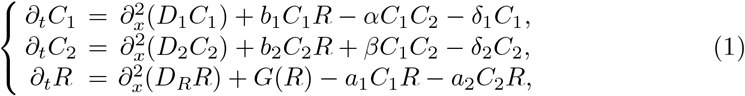

defined on (*t, x*) ∈ (0, *T*) × ℝ_+_. Where *a*_*i*_ is the attack rate of consumer *i* upon the resource, and *b*_*i*_ is the conversion rate of resource into new consumers of species *i* ^3^. The natural mortality of consumer *i* is denoted *δ*_*i*_ and *α* is the attack rate of IG predator upon IG prey, while *β* its conversion rate. The diffusion coefficient of consumer *i* is *D*_*i*_, while the diffusion coefficient of resource is *D*_*R*_.

Finally, in equation (1), the function *G*(*R*) describes how resource grows. We will assume that *G*(*R*) ≥ 0 for 0 ≤ *R ≤ R*^*^ and that *G*(*R*) *<* 0 for *R > R*^*^, such that *G*(*R*^*^) = 0, i.e., resource population grows in the absence of consumers until it attains density *R*^*^. Also, we assume that *R∂*_*R*_*G*(*R*) *< G*(*R*) for *R >* 0, such that resource alongside a single consumer attain stationary states in the model without spatial structure. Throughout the text, we consider the logistic growth function, i.e, *G*(*R*) = *rR*(1 − *R/K*), where *K* is the carrying capacity and *r* the intrinsic growth rate. However, other choices of growth functions that satisfy the conditions stated have similar qualitative results as we will display here, e.g., the chemostat growth function, i.e, *G*(*R*) = *ν* − *rR*, where *ν* is the productivity of the system and *r* resource removing rate, which is often used to model abiotic resources (Klausmeier and Tilman, 2002).

Using the change of variables *τ* = *rt*, 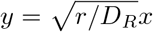, *u*_1_ = (*β/r*)*C*_1_, *u*_2_ = (*α/r*)*C*_2_ and 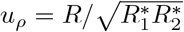, where 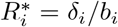 is the resource level under exclusive presence of consumer *i*, we set our equations to

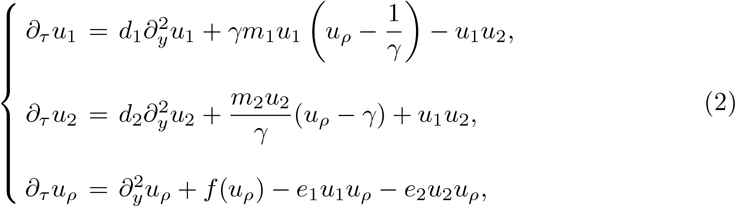

where the new quantities are 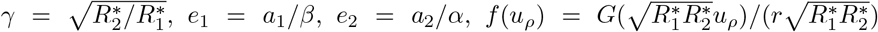, leading to a rescaled carrying capacity 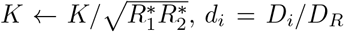 and *m*_*i*_ = *δ*_*i*_*/r*. Coexistence regimes are only possible if *γ >* 1, which is then a competition outcome measure, i.e., whenever *γ >* 1 (*γ <* 1), IG prey (IG predator) is the stronger competitor.

### 2.2 Invasion Regimes and Community formation

Now, we consider Consumer-Resource and IGP communities formed upon introduction of either IG prey or IG predator into a landscape where either resource is established alone or alongside a resident consumer. In order to determine invasibility criteria, we consider a small invading population, such that the equations can be linearized around the invader-free fixed points. In the case of shifts between IG prey and IG predator, such that the former is excluded, the single consumer and resource fixed point is never a center (no sustained oscillations are possible), however, the coexistence fixed point can be either stable or a center of oscillations (see Holt and Polis (1997) for a detailed description)

#### Consumer invades Resource inhabited landscape

To start, we consider a landscape where resource is stablished at 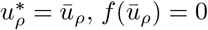, and a small density of IG prey, *u*_1_ ≈ 0, is invading. The leading edge is described by the linearized equation

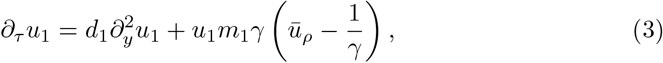

which yields the minimal speed of invasion

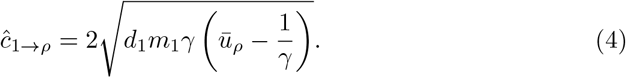

Similarly, IG predator (*u*_2_) invading an resource inhabited landscape will have minimal speed

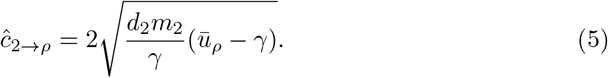

Since a single consumer invading a resource inhabited landscape has linearly determined speed (Lewis et al., 2016; Owen and Lewis, 2001), we have that the asymptotic speeds of invasions equal the minimal ones, i.e., 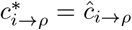. Note that invasion speeds are only real valued for *ū*_*ρ*_ *> γ*^−1^ in the case of IG prey, and *ū*_*ρ*_ *> γ* in the case of IG predator. These set thresholds on parameters of *f* (*u*_*ρ*_). For the logistic growth function, we have *ū*_*ρ*_ = *K*. Then, for *K > K*_1→*ρ*_ = *γ*^−1^ (*K > K*_2→*ρ*_ = *γ*), the landscape can be invaded by IG prey (IG predator).

An example of successful invasion is illustrated in figure 2a (2b) for IG prey (IG predator). Note that as IG prey (IG predator) spreads, the resource level shifts from the carrying capacity *K* to *γ*^−1^ (*γ*). Also, the solutions present the same spatial pattern at different times, i.e., they are traveling wave solutions of the consumer-resource problem, connecting the resource only fixed point at *y* → ∞, to the resource-consumer fixed point at *y* → 0.

**Fig. 2:**
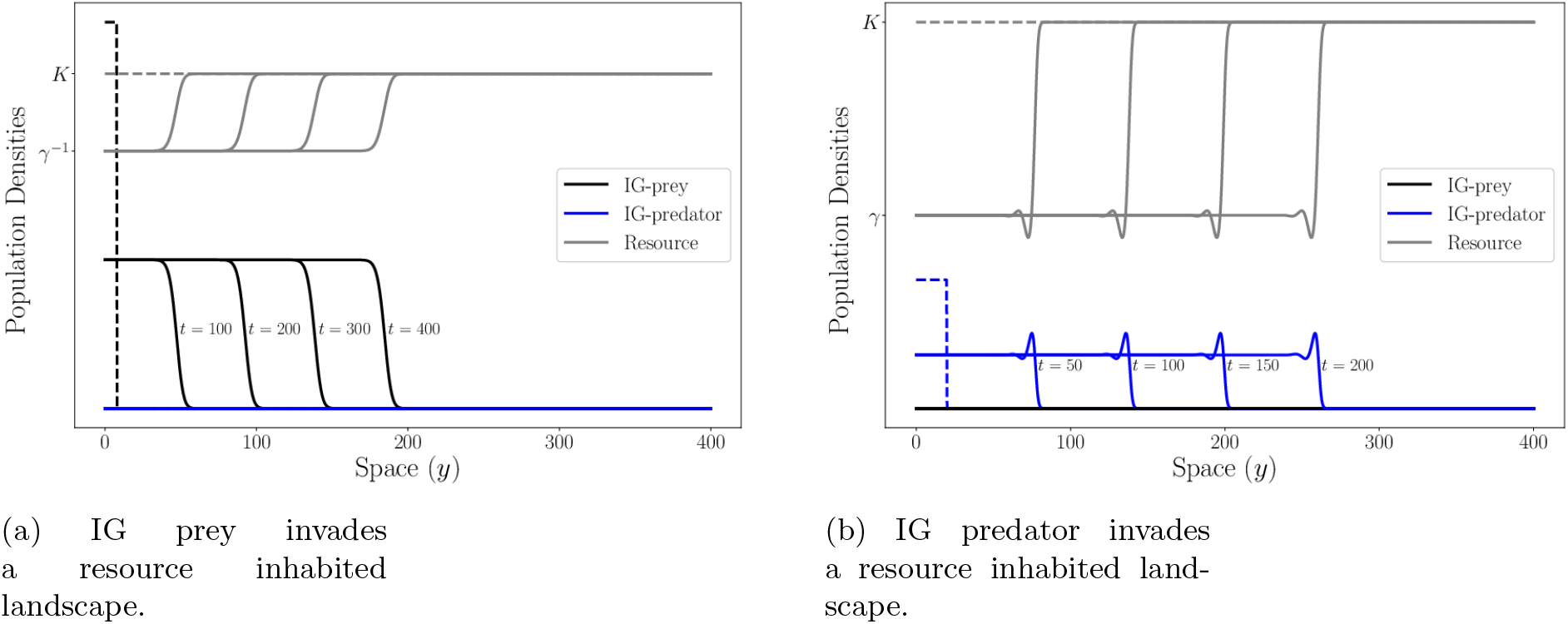
Single consumer invading a resource only inhabited landscape, represented as solutions at different times *t*. Gray lines are Resource, while black (blue) lines are IG prey (IG predator), color matching dashed lines are initial conditions. Parameters are 2*D*_1_ = 2*D*_2_ = *D*_*R*_ = 0.6 (space is not rescaled in the figure), *m*_1_ = 2*m*_2_ = 2*e*_1_ = *e*_2_ = 1.2 and *γ* = 1.5. In (a), *K* = 0.87, while in (b) *K* = 3. We use Neumann boundary conditions on both spatial domain extremities.

#### IG predator invades IG prey and Resource inhabited landscape

When resource is established alongside one of the consumers, however, invasibility criteria change. We start with the case of resource and IG prey stable at levels 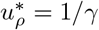 and 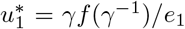, and consider a small density of IG predator invaders, *u*_2_ ≈ 0. The linearized equation is

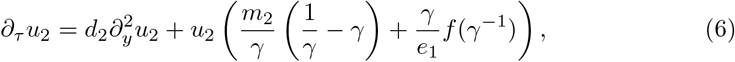

yielding minimal speed

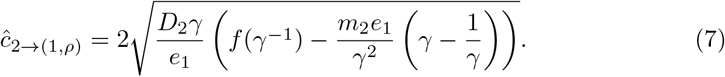

Under the assumption that the minimal speed of invasion is the asymptotic one, IG predator invades a landscape inhabited by resource and IG prey given

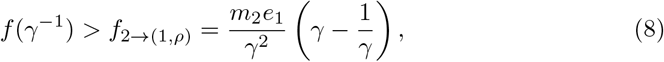

which in turn sets new thresholds for parameter values of *f*. For logistic growth, the carrying capacity must follow

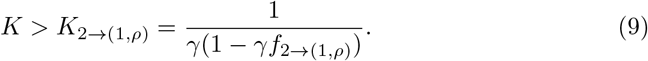

In the case *γ <* 1, condition (8) is always satisfied, so that IG predator always invades and competitively excludes IG prey. For *γ >* 1, in the parameter region *K*_1→*ρ*_ *< K < K*_2→(1,*ρ*)_ we have that only IG prey is able to invade the landscape, while in *K > K*_2→(1,*ρ*)_, IG predator is able to invade the landscape, and, depending on the precise value of *K*, IG prey either coexists alongside IG predator (see figure 3a) or is excluded (figure 3b). Note that, for *γ >* 1, *K*_2→(1,*ρ*)_ *< K*_2→*ρ*_, so the presence of a resident IG prey population facilitates IG predator invasion for these carrying capacity values.

**Fig. 3:**
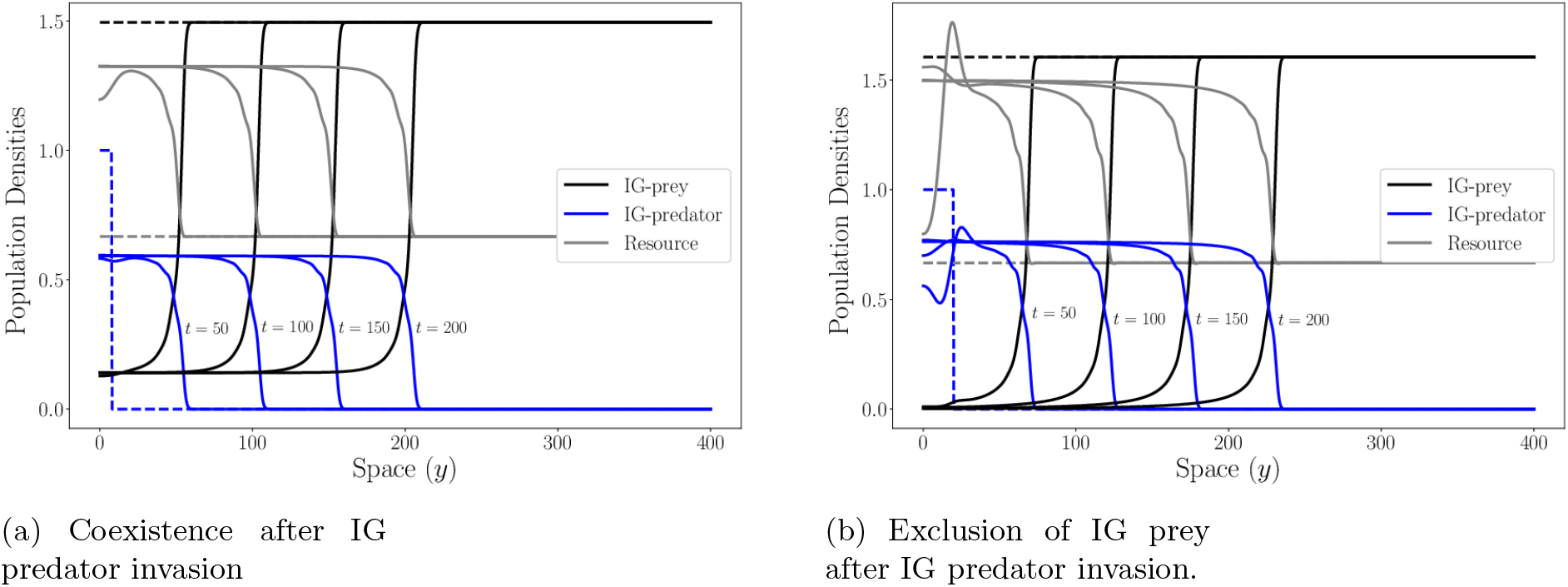
IG predator invading a IG prey and resource inhabited landscape, represented as solutions at different times *t*. Gray lines are Resource, while black (blue) lines are IG prey (IG predator), color matching dashed lines are initial conditions. Parameters are 2*D*_1_ = 2*D*_2_ = *D*_*R*_ = 0.6, *m*_1_ = 2*m*_2_ = 2*e*_1_ = *e*_2_ = 1.2 and *γ* = 1.5. In (a), *K* = 6.5, while in (b) *K* = 18. We use Neumann boundary conditions on both spatial domain extremities and present only the right moving part of the solution.

In figures 3a and 3b we see spatial patterns being formed from *t* = 50 to *t* = 100, and maintained at longer times. In figure 3a, the traveling wave solution connects the IG prey and resource fixed point at *y* → ∞ to the coexistence one in *y* → 0, while in figure 3b, it connects the IG prey and resource fixed point to the IG predator and resource fixed point.

#### IG prey invades IG predator and Resource inhabited landscape

Finally, we consider a landscape inhabited by resource and IG predator, at stable densities 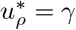 and 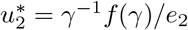. A small density of IG prey, *u*_1_ ≈ 0, is described by the linearized equation

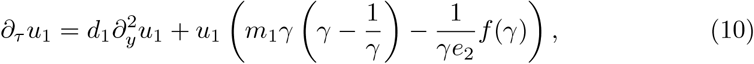

and has minimal speed

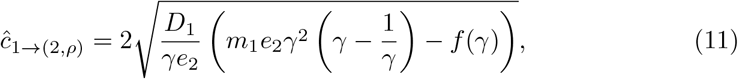

yielding yet another threshold for *f* in which IG prey is able to invade the landscape, given by

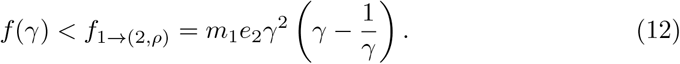

The threshold for carrying capacity is

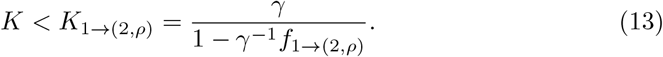

For *γ <* 1, i.e., when IG prey is not the best competitor, its minimal invasion speed is never real valued. Assuming that the minimal speed is the asymptotic one, we have that IG prey is never able to invade. For *γ >* 1, *K*_2→(1,*ρ*)_ *< K < K*_1→(2,*ρ*)_ is a mutual invasibility region, i.e., IG prey can invade an IG predator occupied landscape and vice-versa, leading to a region of coexistence between both consumers (see figure 4). The region *K > K*_1→(2,*ρ*)_ IG prey can no longer invade an IG predator occupied landscape, and, in turn, IG predator invasions lead to IG prey exclusion, as previously shown in figure 3b. Also for *γ >* 1, we have *K*_2→(1,*ρ*)_ *< K*_2→*ρ*_, and IG prey can never competitively exclude a resident IG predator population uppon invasion.

**Fig. 4:**
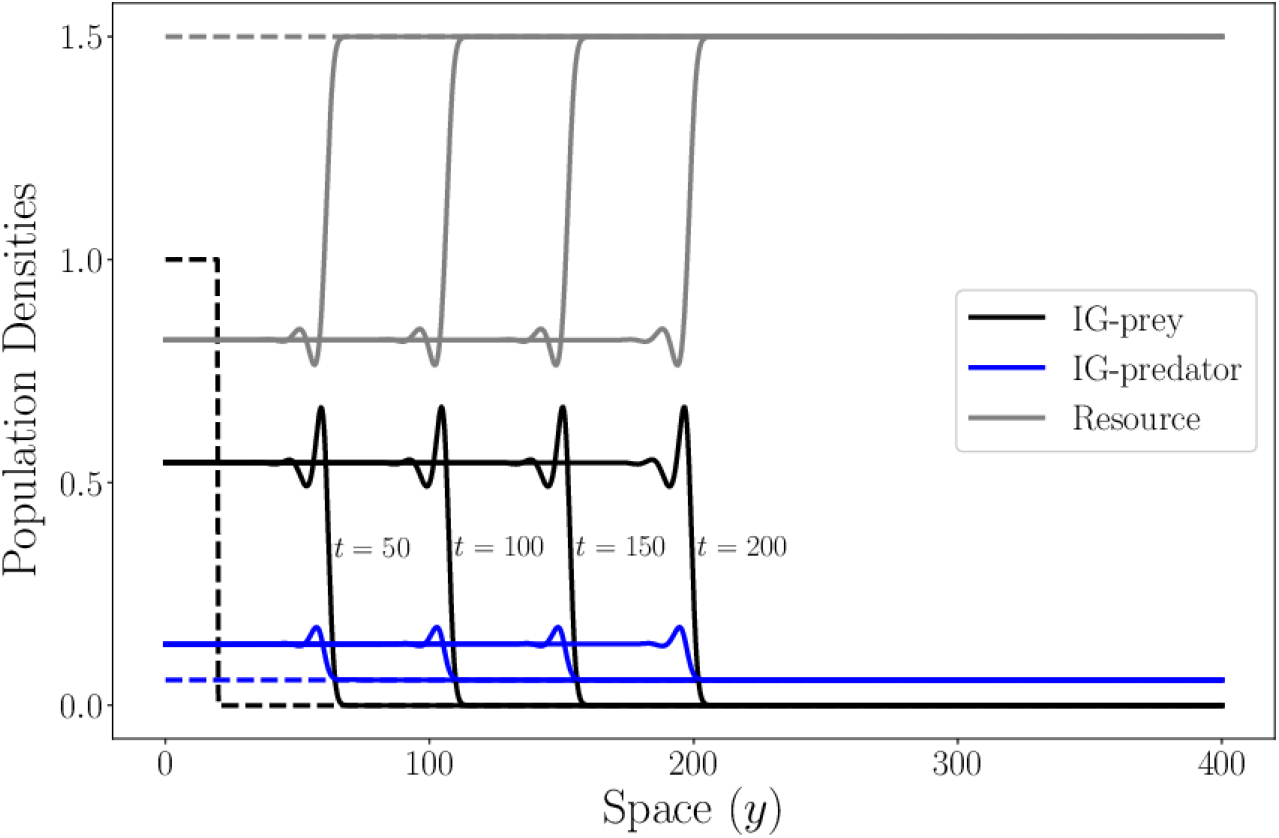
IG prey invading a IG predator and resource inhabited landscape, represented as solutions at different times *t*. Gray lines are Resource, while black (blue) lines are IG prey (IG predator), color matching dashed lines are initial conditions. Parameters are 2*D*_1_ = 2*D*_2_ = *D*_*R*_ = 0.6, *m*_1_ = 2*m*_2_ = 2*e*_1_ = *e*_2_ = 1.2 and *γ* = 1.5, *K* = 1.6. Traveling wave solutions connect the resident IG predator and resource fixed point to the coexistence one. We use Neumann boundary conditions on both spatial domain extremities and present only the right moving part of the solution.

When the coexistence fixed point is unstable and display maintained oscillations, the system can display dynamical stabilization (Malchow and Petrovskii, 2002; Petrovskii and Malchow, 2000). In figure 5, as the front of invasion advances with speed *c*_*i*→(*j,ρ*)_, the interface between the dynamical stability region and oscillatory regime at the core of invasion also advances in space, but with a smaller speed than the front, such that the length of the dynamical stability region is increasing throughout invasion, similar to what is found in Malchow and Petrovskii (2002).

**Fig. 5:**
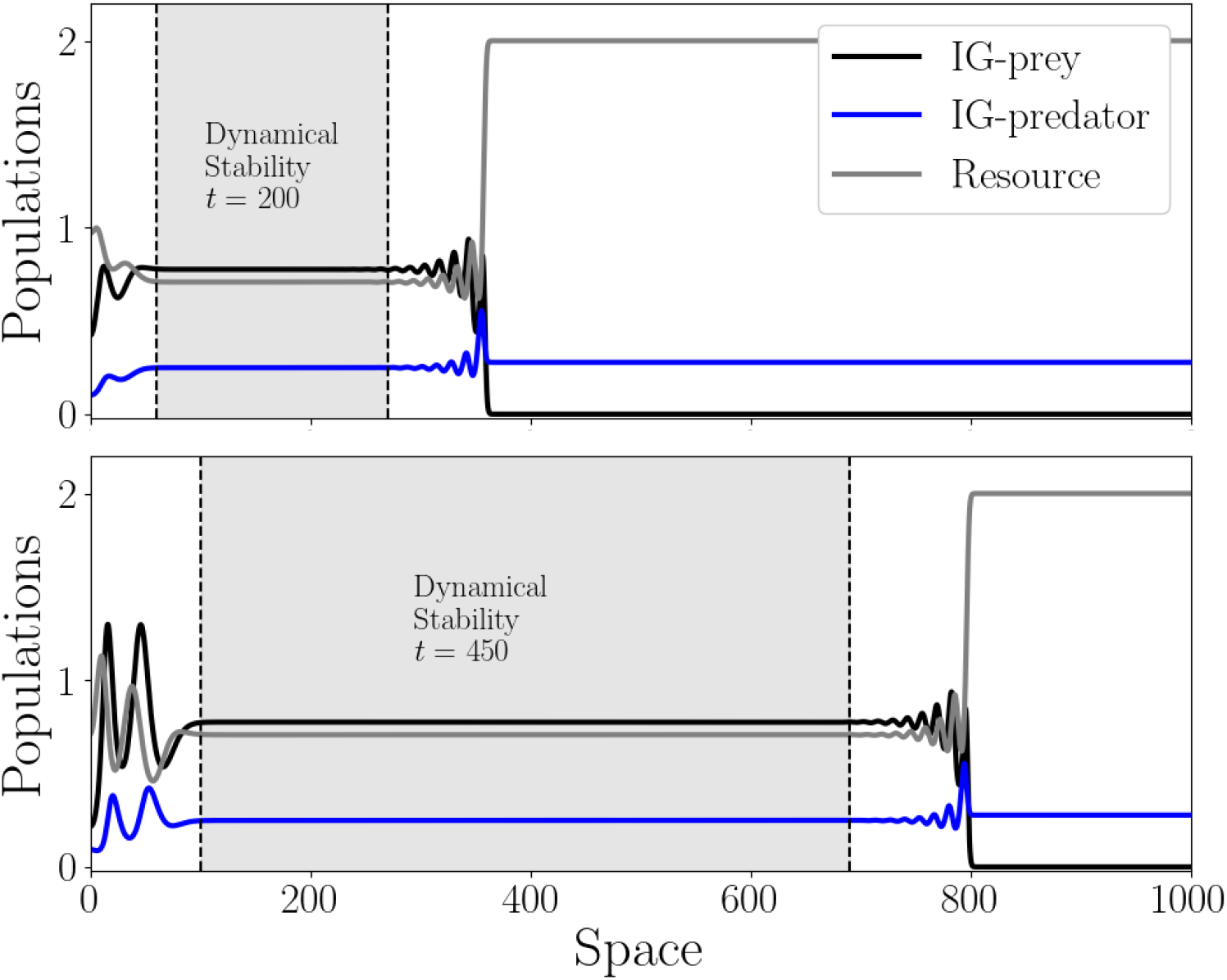
IG prey invasion leading to dynamical stability (light gray colored region) at *t* = 200 (top) and *t* = 450 (bottom. Parameters used are *K* = 3, *γ* = 2 and *e*_2_ = 2*e*_1_ = *m*_1_ = 2*m*_2_ = 1.2, *D*_1_ = *D*_2_ = *D*_*R*_ = 0.5

### 2.3 Asymptotic Invasion Speeds

We assumed that the minimal invasion speeds correspond to the asymptotic ones. Measuring the invasion speeds numerically (dots in figures 6a-6d) reveals that, for the explored region of parameter space, this is indeed the case, i.e., the asymptotic invasion speeds equal the minimal ones (solid lines in figures 6a-6d).

**Fig. 6:**
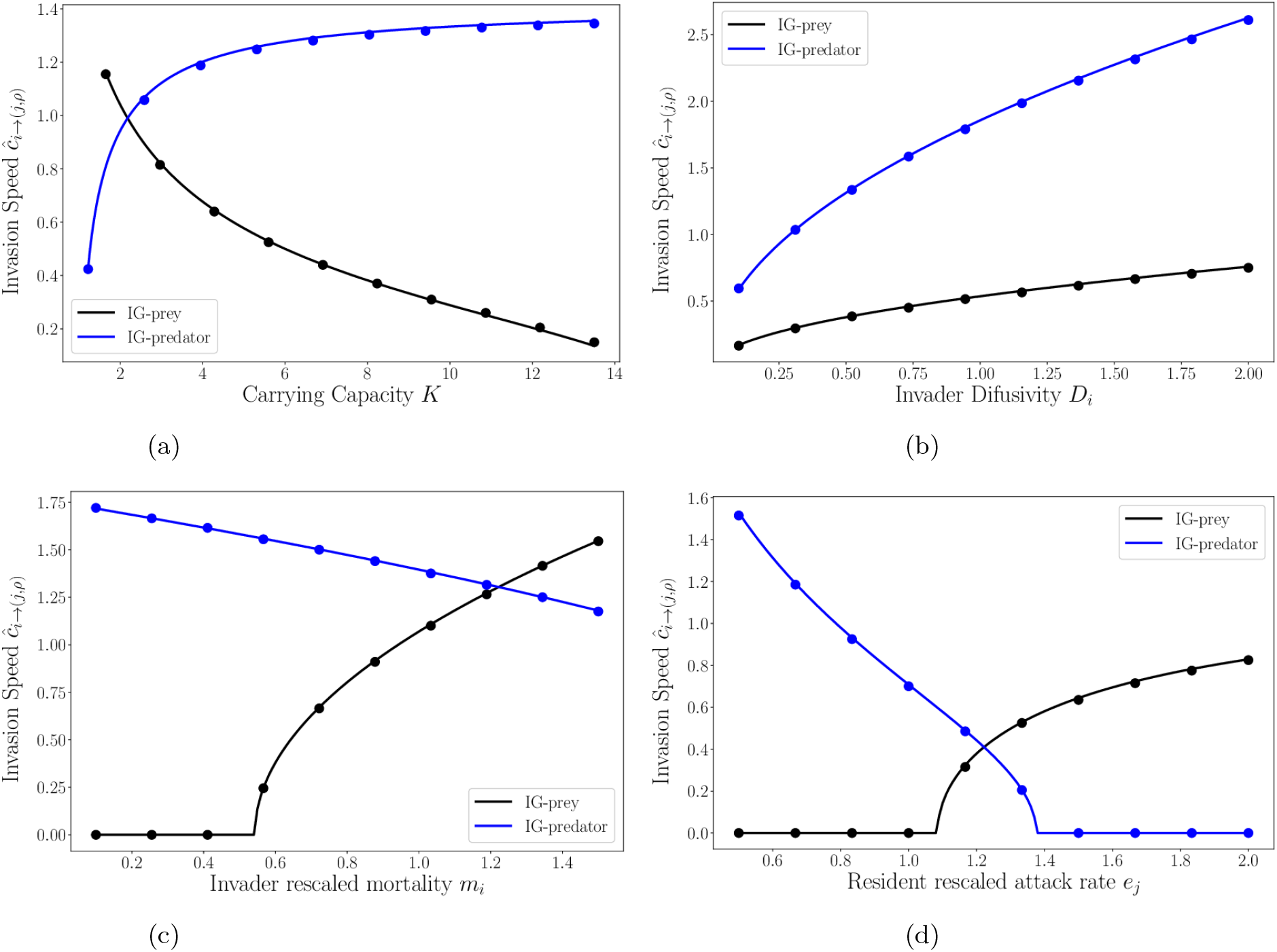
Numerical (dots) and linearly obtained (lines) invasion speeds for IG prey (black) and IG predator (blue). We always analyze the cases of invasion uppon a consumer-resource community. We have speed as a function of: carrying capacity in panel 6a, diffusivity in panel 6b, mortality in 6c and attack rate in 6d. Parameters used, with the exception of the varying ones on the *x* axis in each of their respective figures, are *D*_1_ = *D*_2_ = *D*_*R*_ = 0.5, *γ* = 1.5, *m*_2_ = 2*m*_1_ = 2*e*_1_ = *e*_2_ = 1.2, *K* = 3

Note that increasing carrying capacity (figure 6a) increases IG predator invasion speed while decreases IG prey invasion speed, as expected, since we have a lower threshold for IG predator invasion in terms of carrying capacity (*K*_2→(1,*ρ*)_) and an upper treshold for IG prey invasion (*K*_1→(2,*ρ*)_). Increasing the invading species diffusivity (figure 6b) increases both consumers invasion speeds, as expected.

Invasion speeds behavior in respect to *m*_*i*_ and *e*_*i*_ is not that trivial. Note that increasing rescaled mortalities *m*_*i*_ and resident consumer attack rates upon resource *e*_*j*_ produce opposite behaviors on IG prey and IG predator. The main reason lies in the fact that, at the leading edge, the net effect of competition for resource is positive (negative) in IG prey (IG predator). Since the net effect of competition in consumer *i* is proportional to *m*_*i*_, increases in *m*_1_ increase IG prey invasion speed, while the opposite holds for IG predator. A similar discussion can be made in terms of *e*_*i*_ and the net effect of intraguild predation at the leading edge. While increasing *e*_1_ decreases the total available IG prey for the consumption of an invading IG predator, thus, reducing its invasion speed, increasing *e*_2_ decreases the total amount of IG predator, reducing predation pressure on an invading IG prey and allowing it to spread with faster speeds.

To illustrate the effects of invasion leading to both regimes on the spreading speeds, we calculate numerical spreading speeds (dots in figure 7) and compare them to the ones obtained from linearization (solid lines in figure 7) in a parameter region where the coexistence fixed point is unstable (region *γ >* 2 in figure 7). We observe that, although traveling wave solutions are not being formed, the spreading speeds are still linearly determined and expressions (7) and (11) provide accurate estimates. This was expected, since it also holds in the case of dynamical stabilization regimes of predator-prey models (Malchow and Petrovskii, 2002).

**Fig. 7:**
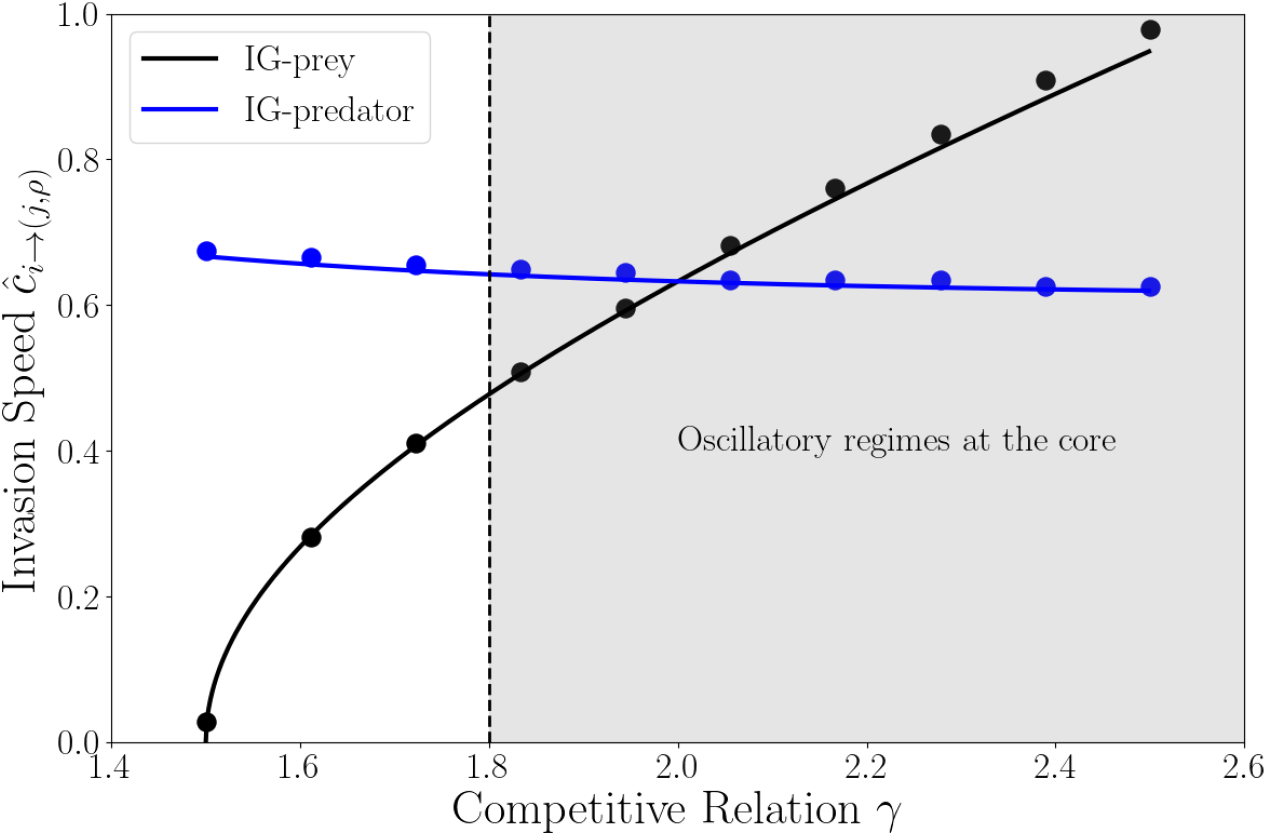
Assymptotic Invasion Speeds for different values of *γ*. In the region *γ >* 1.8 (roughly) dynamic stabilization regimes are possible. The numerically obtained speed (dots) match expressions (7) and (11) (lines). Parameters used are *K* = 5 and *e*_2_ = 2*e*_1_ = *m*_1_ = 2*m*_2_ = 1.2, *D*_1_ = *D*_2_ = *D*_*R*_ = 0.5

## 3 Intraguild Predation in Heterogeneous Landscapes

### 3.1 Model

We follow Maciel and Lutscher (2013); Yurk and Cobbold (2018); Cobbold et al. (2022) closely. We let the space be composed of two types of patches, 1 and 2, of sizes *l*_1_ and *l*_2_ respectively, displaced periodically on the real line. We denote the desities of IG prey inside patches of the *j*-th type as *C*_1*j*_, while for IG predators and Resource we use *C*_2*j*_ and *R*_*j*_, respectively. The dynamics of these populations on a patch of type *j* are given by

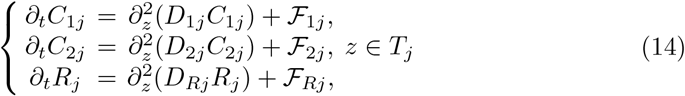

where *T*_*j*_ is the set of points within patches of type *j*, i.e.,

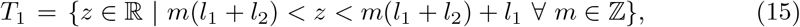

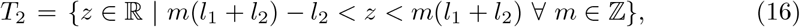

and ℱ_*ij*_ ≡ ℱ_*ij*_(*C*_1*j*_, *C*_2*j*_, *R*_*j*_), *i* = 1, 2, *R*, are the growth functions of species *i* at a patch of type *j*. Following our model in homogeneous space 1 and letting intraguild predation relations occur in both patches we have

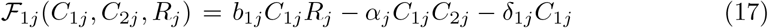

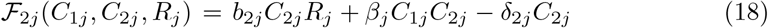

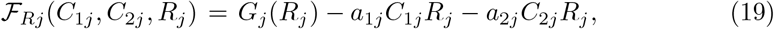

with symbols maintaining their definition as in (1), but now containing an extra index, *j*, to denote the patch type in which they are valid. The same is true for the difusion coeficcients *D*_*ij*_. Also, we keep *G*_*j*_, *j* = 1, 2, as a logistic growth function ^4^, with intrinsic growth rate *r*_*j*_ and carrying capacity *K*_*j*_. For the description of parameters and their correspondence to the homogeneous model (1), check table 1.

**Table 1:**
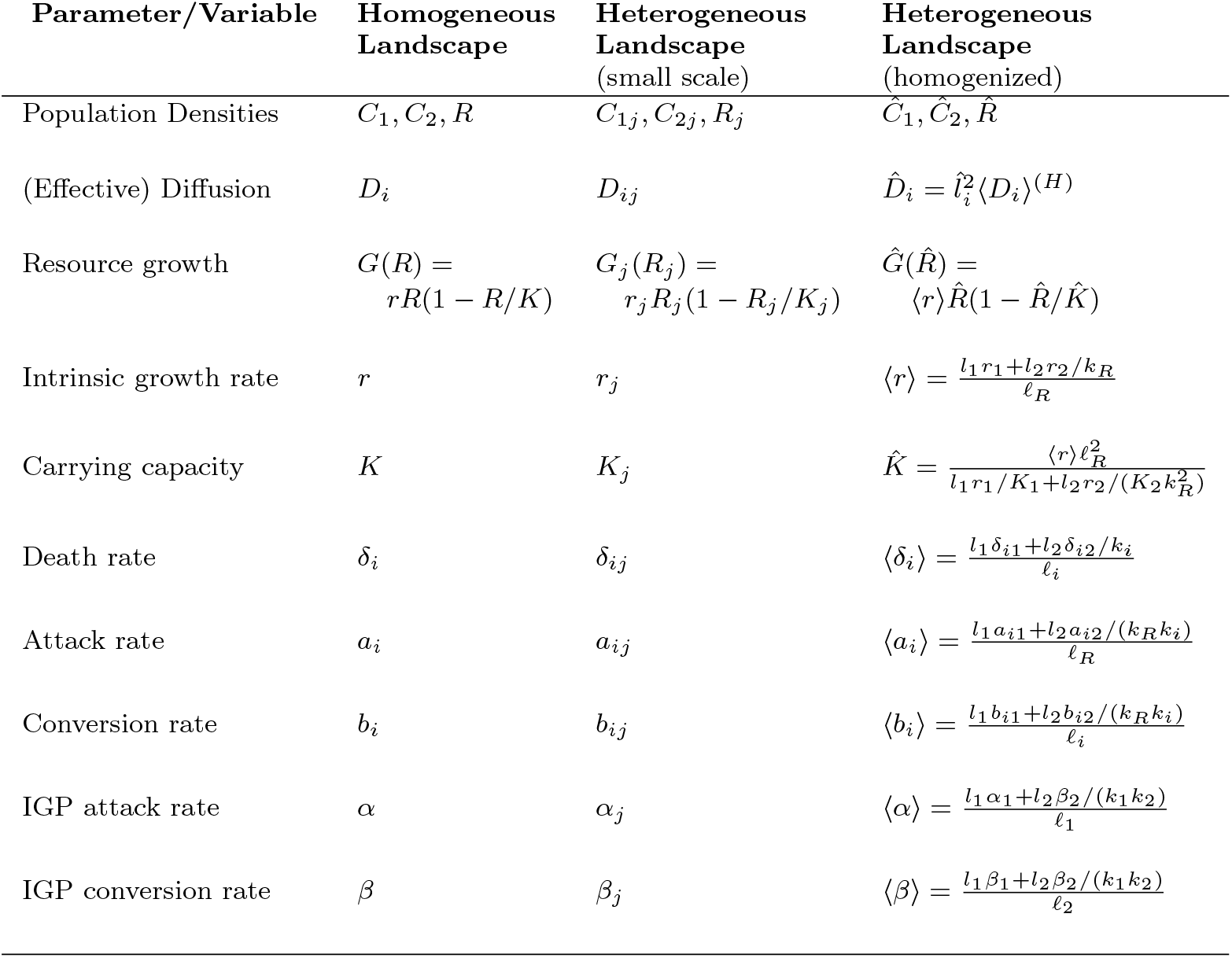
Symbol correspondence for parameters on models (1), (14) and (25).

At the interface *z*_*m*_ of patches of type 1 and 2, we assume continuous flux, but discontinuous densities, to account for habitat preference (Ovaskainen and Cornell, 2003; Maciel and Lutscher, 2013), leading to

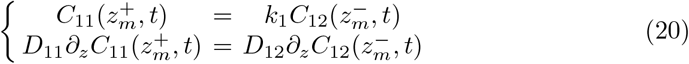

where *z*_*n*_ = *n*(*l*_1_ + *l*_2_) + *ζ*_*n*_*l*_1_, *ζ*_*n*_ = 1 (*ζ*_*n*_ = 0) if *n* is odd (even), and *k*_1_ is the IG prey density effective patch preference. We proceed similarly for *C*_2_ and *R* to write their interface conditions and define the patch preferences, *k*_2_ and *k*_*R*_, for the IG predator and the resource, respectively.

We follow Maciel and Lutscher (2013) to set

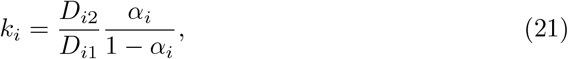

where *α*_*i*_ ∈ (0, 1) is the probability of species *i, i* = 1, 2, *R*, to move from the interface into a type 1 patch.

It is helpful to write the model (14) in a shorter notation. We define the piece-wise constant (in *z*) functions *D*_*i*_(*z*) = *D*_*ij*_, *z* ∈ *T*_*j*_ and ℱ_*i*_(*z*, ·) = ℱ_*ij*_(·), *z* ∈ *T*_*j*_, to write

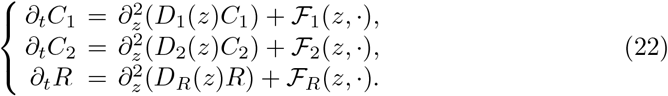

Of course, (22) is only equivalent to (14) when interface conditions (20) are accounted for. However, this notation allows us to quickly address population densities *C*_1_, *C*_2_ and *R* in the landscape level (across multiple different patches).

### 3.2 Homogenization Technique

We proceed with a multiscale analysis and approximation method following Yurk and Cobbold (2018), we will briefly outline the method, but refer to the original study for more details. Also, we will describe the methods in terms of a single species, *C*_1_, but the proceedings are the same for *C*_2_ and *R*, and are to be taken simultaneously.

We define the large scale *x* and the small scale *z* = *x/*𝓁, with 𝓁 = *l*_1_ + *l*_2_ ≪ 1 in the large scale, and assume that population densities depend on both *x* and *z*, leading to *C*_1_ ≡ *C*_1_(*x, z, t*), the IG prey density in both. Expanding such solutions in 𝓁, we get

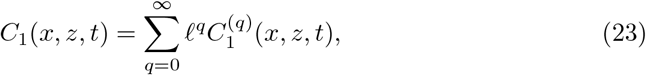

and assuming *x* and *z* are independent we have 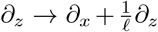. When substituting the expanded solution and the change of variables to (14) with interface conditions (20), we find a system of coupled equations, that can be solved for 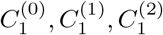.

The leading order of the expansion, 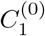, is given by

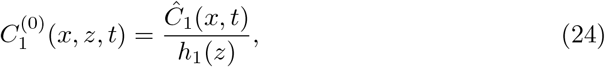

where *h*_1_(*z*) = 1 (*h*_1_(*z*) = *k*_1_) for *z* ∈ *T*_1_ (*z* ∈ *T*_2_). Applying the same procedure to *C*_2_ and *R*, we arrive in a similar expression for 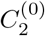 and *R*^(0)^, with *h*_2_(*z*) and *h*_*R*_(*z*) defined in the same fashion as *h*_1_(*z*). Since *h*_*i*_(*z*) *>* 0 ∀*i, z*, the population densities are strongly dependent on *Ĉ*_1_, *Ĉ*_2_ and 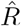, which are obtained by solving

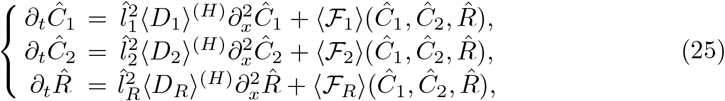

where, 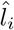, ⟨*D*_*i*_⟩^(*H*)^ and ⟨ℱ⟩_*i*_, are the scaled spatial periods, diffusion coefficients and growth functions of species *i* in the large scale, respectively. Namely, the scaled periods are given by

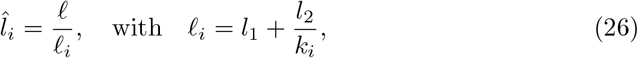

while the diffusion coefficients are the harmonic mean between the diffusion coefficients of each habitat, i.e.,

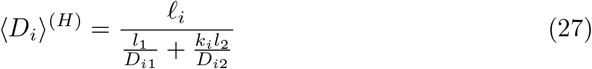

and the growth functions are the arithmetic mean of ℱ_*ij*_, i.e.,

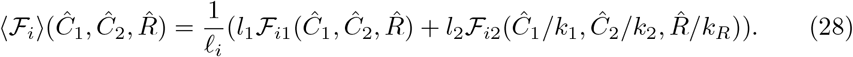

Rearranging the terms in ⟨ℱ_*j*_ ⟩, we find that system (25) can be written in the same form as (1), i.e.

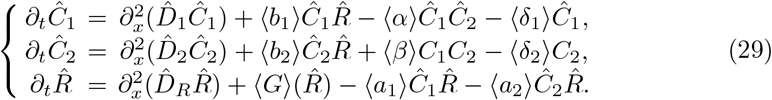

We organize the definition and correspondence between the heterogeneous landscape homogenized parameters in model (29) and the homogeneous landscape parameters in model (1) in table 1.

With the parameters defined in table 1, we rescale time, space and population densities as follows:

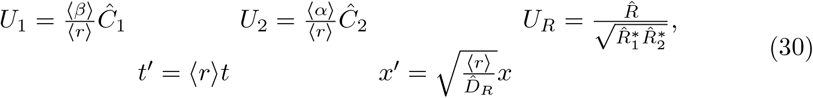

where

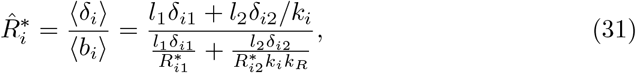

is the (approximate) resource level when established with only consumer *i* in an heterogeneous landscape. 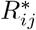 has a similar definition as 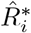 (and 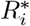 in (1)), but only with respect to the patch type *j*, i.e., 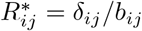, *j* = 1, 2 and *i* = 1, 2.

After some algebra, system (29) can be written as

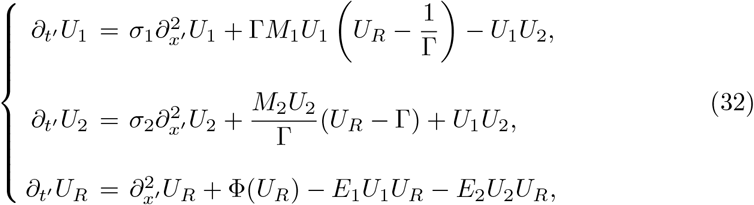

which is precisely in the same form of equation (2). The new quantities are

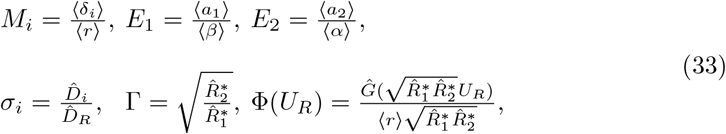

where we rescale the carrying capacity to 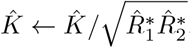.

### 3.3 Mutual Invasibility Conditions

Since models (32) and (2) are in the same form, we expect mutual invasibility of IG prey and IG predator to take place in the same correspondent parameter regions as found for the homogeneous landscapes. This way, we expect mutual invasibility only for Γ *>* 1, i.e., when the homogenized competitive measure shows that IG prey is the stronger competitor, and within a range of effective carrying capacities 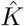.

The condition Γ *>* 1, in full form, becomes

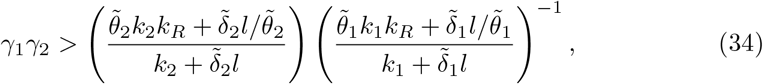

where *l* = *l*_2_*/l*_1_ is the ratio of patch sizes, 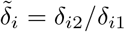 is the ratio of species *i* death rates in patch 2 and patch 1, 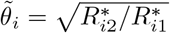, the ratio between resource levels in presence of species *i* in different patch types. Finally, 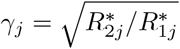 is the competition outcome measure in patches of type *j*.

Whenever the right-hand side of inequality (34) is smaller than unity, mutual invasibility is facilitated and can occur even if *γ*_*j*_ *<* 1, *j* = 1, 2, i.e., competitive reversals in favor of IG prey are possible. At the same time, whenever the right-hand side is larger than unity, mutual invasibility is hindered, and may not occur even if *γ*_*j*_ *>* 1, *j* = 1, 2, i.e., competitive reversals in favor of IG predator are also possible. When the right-hand side is precisely unity, the condition reduces to *γ*_1_ *> γ*^−1^, that is, whenever IG prey is not competitively stronger in one of the patches, it has to overcompensate this effect in the other patch.

The possible competitive reversal scenarios found on inequality (34) depend heavily on all three species patch preferences, *k*_*i*_, *i* = 1, 2, *R*, how patchy the landscape is, given by *l*, as well as consumer traits (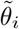 and 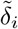). To simplify this relation and gain some insight on how habitat preferences govern this inequality, we assume *γ*_1_ = *γ*_2_ = *γ*, which also implies 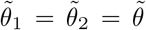, and define 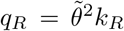 and 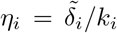. With this, condition (34) becomes

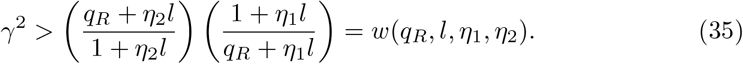

Note that *w*(*q*_*R*_ = 1, ·) = 1, i.e., whenever *q*_*R*_ = 1, we recover the conditions found in an homogeneous landscape. Moreover,

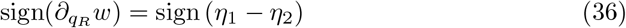

such that if *η*_1_ *> η*_2_, then *w* is monotonically increasing in *q*_*R*_, and monotonically decreasing otherwise. Because *q*_*R*_ is monotone in 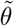 and *k*_*R*_, *w* is also monotone in these parameters. Since *w*(*q*_*R*_ = 1, ·) = 1, if *w* is monotonically increasing in *q*_*R*_ (figure 8a), then for *q*_*R*_ *>* 1 we have hindered mutual invasibility conditions, because *γ*^2^ *> w*(*q*_*R*_, ·) *>* 1, i.e., *γ* must be larger than what is expected in an homogeneous landscape. For *q*_*R*_ *<* 1 we have facilitated mutual invasibility conditions, because *γ*^2^ *> w*(*q*_*R*_, ·) with *w*(*q*_*R*_, ·) *<* 1, i.e., *γ* can be smaller than what is expected in an homogeneous landscape. The opposite holds when *w* is monotonically decreasing (figure 8b).

**Fig. 8:**
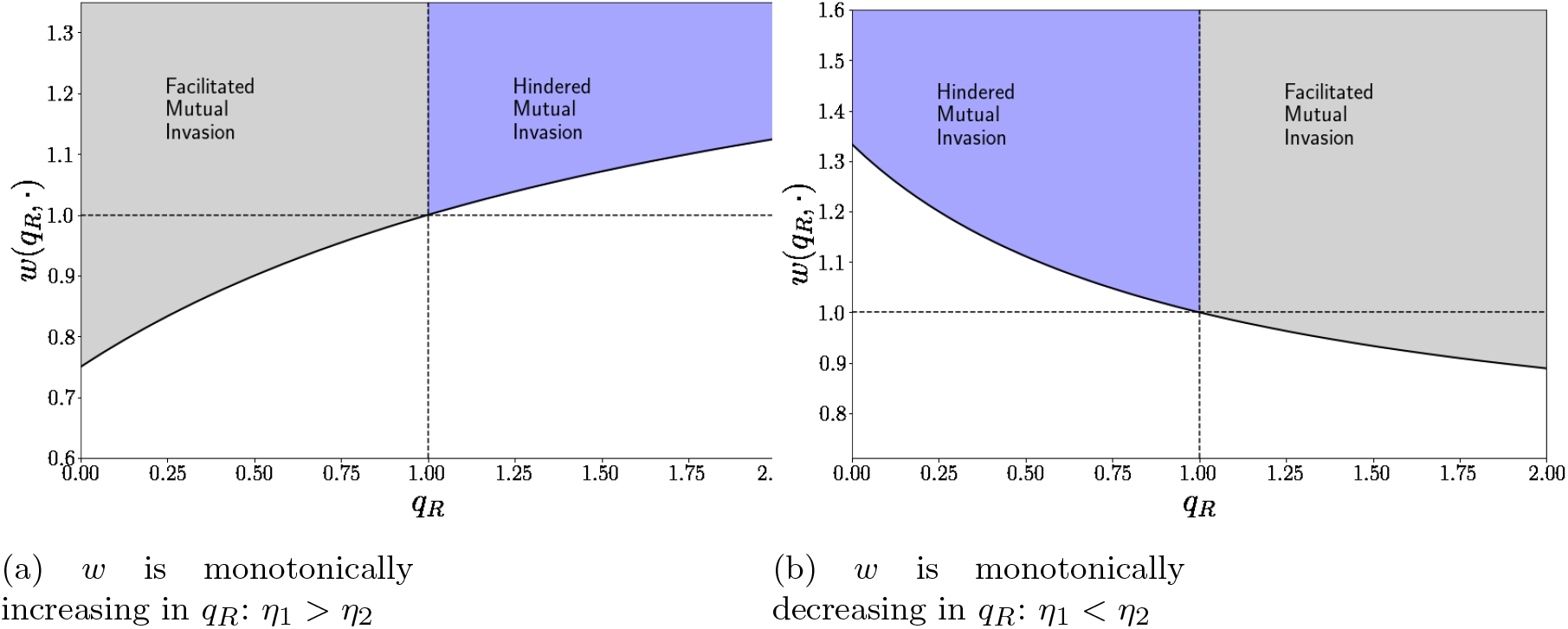
Regimes of facilitated and hindered mutual invasions, depending on how the quantities 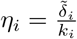, *i* = 1, 2 relate.

The ecological interpretation of inequality (35) and equation (36) is as follows: 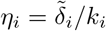 relates species *i* patch preference, *k*_*i*_, and how much larger is its death rate in pathces of type 2 compared to type 1, 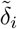. Whenever there is a difference between IG prey and IG predator in this relation, then mutual invasibility is hindered or facilitated depending on how unbalanced is resource consumption between different patches, *θ*, and resource patch preference *k*_*R*_.

To understand solely the effects of patch preferences, let 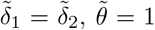 and con-sider the following example: IG prey prefers patches of type 1, such that *k*_1_ *>* 1, and IG predator prefers patches of type 2, such that *k*_2_ *<* 1, this way, *w* is monotonically decreasing. The region *k*_*R*_ *>* 1 (resource prefers patches of type 1) leads to a competitive reversal in favor of IG prey whenever *w < γ*^2^ *<* 1, while the region *k*_*R*_ *<* 1 (resource prefers patches of type 2) leads to competitive reversals in favor of IG predator whenever *w > γ*^2^ *>* 1. The outcomes of the example are reversed when *k*_1_ *<* 1 *< k*_2_, i.e., IG prey and IG predator patch preferences are switched. This way, whichever consumer has their patch preferences aligned with resource species’ patch preference can benefit from competitive reversals.

Similarly, we can understand competitive reversal scenarios only considering the death rate ratios, 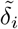, and uneven resource consumption between patches, 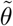, through the following example: Let *k*_1_ = *k*_2_ and *k*_*R*_ = 1 and consider IG prey have a smaller death rate in patches of type 1 than in patches of type 2, and the opposite for IG predator, such that 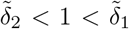. With that, *w* is monotonically increasing in 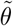. For 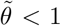, i.e., resource is less consumed in patches of type 1, we may have competitive reversals in favor of IG prey whenever *w < γ*^2^ *<* 1, while 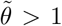 favors IG predator if *w > γ*^2^ *>* 1. That way, whenever the patch where IG prey dies less is also the one where resource is less consumed, we have facilitated mutual invasibility, while hindered conditions apply when we have the opposite.

We could also discuss similar effects that occur between *k*_1_, *k*_2_ and 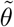, as well as between 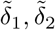 and *k*_*R*_, but the outcome would be quite similar. In a general manner, whenever the conditions favor IG prey (IG predator) in some way, the mutual invasibility conditions are facilitated (hindered). To formally write these parameter regions, we define *δη* = *η*_1_ −*η*_2_ and write the facilitated, ℋ_*f*_, and hindered, ℋ_*h*_, mutual invasion conditions as quadrants in the parameter space (*q*_*R*_, *δη*) ∈ ℝ_+_ × ℝ defined by

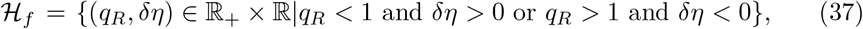

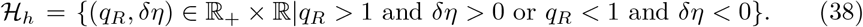

The ratio among patch sizes, *l*, does not appear in any of the relations discussed so far, but it does play an important role. Again, we focus on the case *γ*_1_ = *γ*_2_ = *γ*, just to simplify expressions and gain some insight.

We have Γ^2^ = *γ*^2^*/w*, and we note that

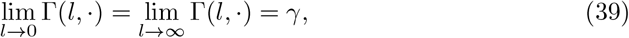

i.e., whenever the landscape is almost homogeneous (*l*_1_ ≫ *l*_2_ or *l*_2_ ≪ *l*_1_), we recover the homogeneous mutual invasion conditions.

A quick inspection on the derivative *∂*_*l*_Γ^2^ reveals that

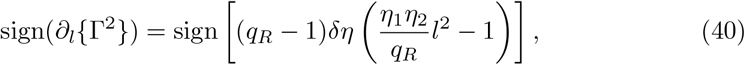

so *∂*_*l*_Γ switches sign just once, at 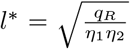, which is therefore the only extremum point of Γ w.r.t *l*, and limits (39) imply Γ is bounded by its extremum, Γ(*l* = *l*^*^, ·), and *γ*. Note that *l*^*^ is the maximum point of Γ(*l*, ·) whenever (*q*_*r*_, *δη*) ∈ ℋ_*f*_, and by limits in (39) we have *γ <* Γ(*l*, ·) ≤ Γ(*l* = *l*^*^, ·). Therefore, if *γ >* 1, mutual invasion regimes are possible for any *l*. In a similar fashion, *l*^*^ is a minimum point of Γ(*l*, ·) whenever (*q*_*r*_, *δη*) ∈ ℋ_*h*_, and the limits in (39) imply Γ(*l* = *l*^*^, ·) ≤ Γ(*l*, ·) *< γ*. Therefore, if *γ <* 1 mutual invasion regimes are not possible for any *l*. This reads that whenever IG prey (resp. IG predator) is the superior competitior and is benefited by facilitated (resp. hindered) mutual invasion conditions, mutual invasions can (resp. do not) take place regardless of how patchy the landscape is.

Now, assuming that *l*^*^ is a maximum point, in order to have Γ(*l* = *l*^*^, ·) *>* 1, we must have

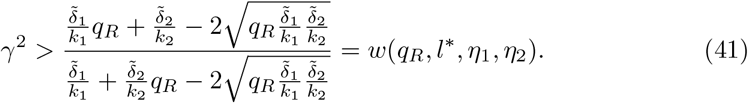

However, *w*(*q*_*R*_, *l*^*^, *η*_1_, *η*_2_) *<* 1 for (*q*_*R*_, *δη*) ∈ ℋ_*f*_. By continuity of Γ(*l*, ·), whenever

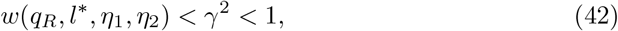

we have competitive reversals in favor of IG prey around a neighborhood of *l*^*^ and mutual invasions are possible, i.e., in order to have Γ *>* 1 even when *γ <* 1, the proportion of patch type lengths must be close to *l*^*^.

The precise extent of *l* values at which competitive reversals occur is obtained by solving 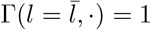, which yields 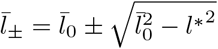, where

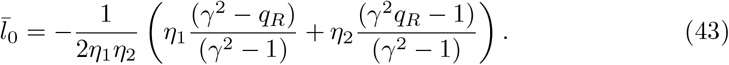

Note that roots 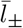 become negative whenever *γ >* 1 and (*q*_*R*_, *δη*) ∈ ℋ_*f*_. In this regime mutual invasion become possible for any *l*, as expected.

By assuming that *l*^*^ is a minimum, i.e., that hindered mutual invasion conditions take place, we arrive at complementary results. We have

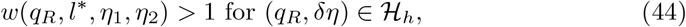

and whenever

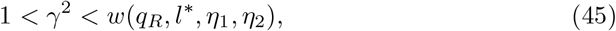

competitive reversals occur in favor of IG predator. Mutual invasions are only possible outside the rang 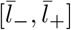, i.e., when the landscape is more homogeneous/less patchy. Also, if *γ <* 1 and (*q*_*R*_, *δη*) ∈ ℋ_*h*_, both 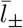 become negative and mutual invasibility is not possible, as expected.

Whenever competitive reversals occur, (*q*_*R*_, *δη*) determines in which direction the reversals are and the roots *l*_*±*_ delimit the heterogeneity levels of the landscape necessary for it. Still, mutual invasibility regions only take place within a range of effective carrying capacities, 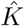. We proceed as in the homogeneous case and define

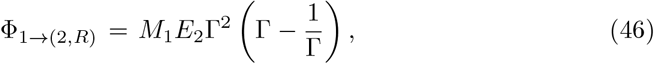

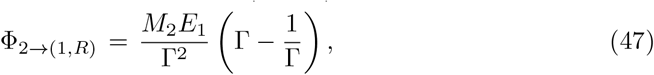

and with these, the threshold values in 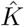 become

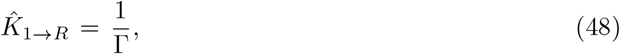

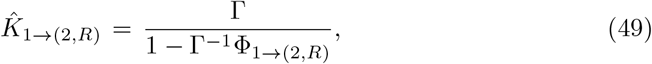

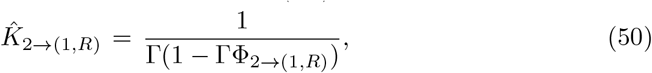

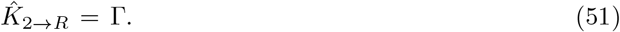

The mutual invasibility region is delimited by 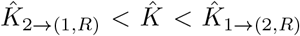, which only exists if Γ *>* 1. We investigate these regions in the plane 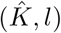 by plotting the different threshold values 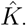. To illustrate the precise effects of competitive reversals, we focus again on *γ*_1_ = *γ*_2_ = *γ*.

First, let us consider the case of competitive reversals in favor of IG prey, we consider *w*(*q*_*R*_, *l*\*, *η*_1_, *η*_2_) *< γ <* 1 and (*q*_*R*_, *δη*) ∈ ℋ_*f*_, to plot figure 9. At the limits log(*l*) → ±∞, the landscape is more homogeneous and heavily composed of a single patch type, and regime shifts from increasing levels of carrying capacity occur only from resource alone (pastel colored) to IG predator and resource communities (blue colored). Note that the 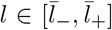 region, when the landscape become more patchy, we have Γ *>* 1 in the upper plot, corresponding to the *l* region where IG prey can establish alone with resource (gray colored) and mutual invasions can occur (cyan colored) in the lower plot. All curves are increasing in log(*l*) because *k*_*R*_ *<* 1*/*2, so resource strongly prefers patches of type 1, which length’s proportion decrease with increasing *l*.

**Fig. 9:**
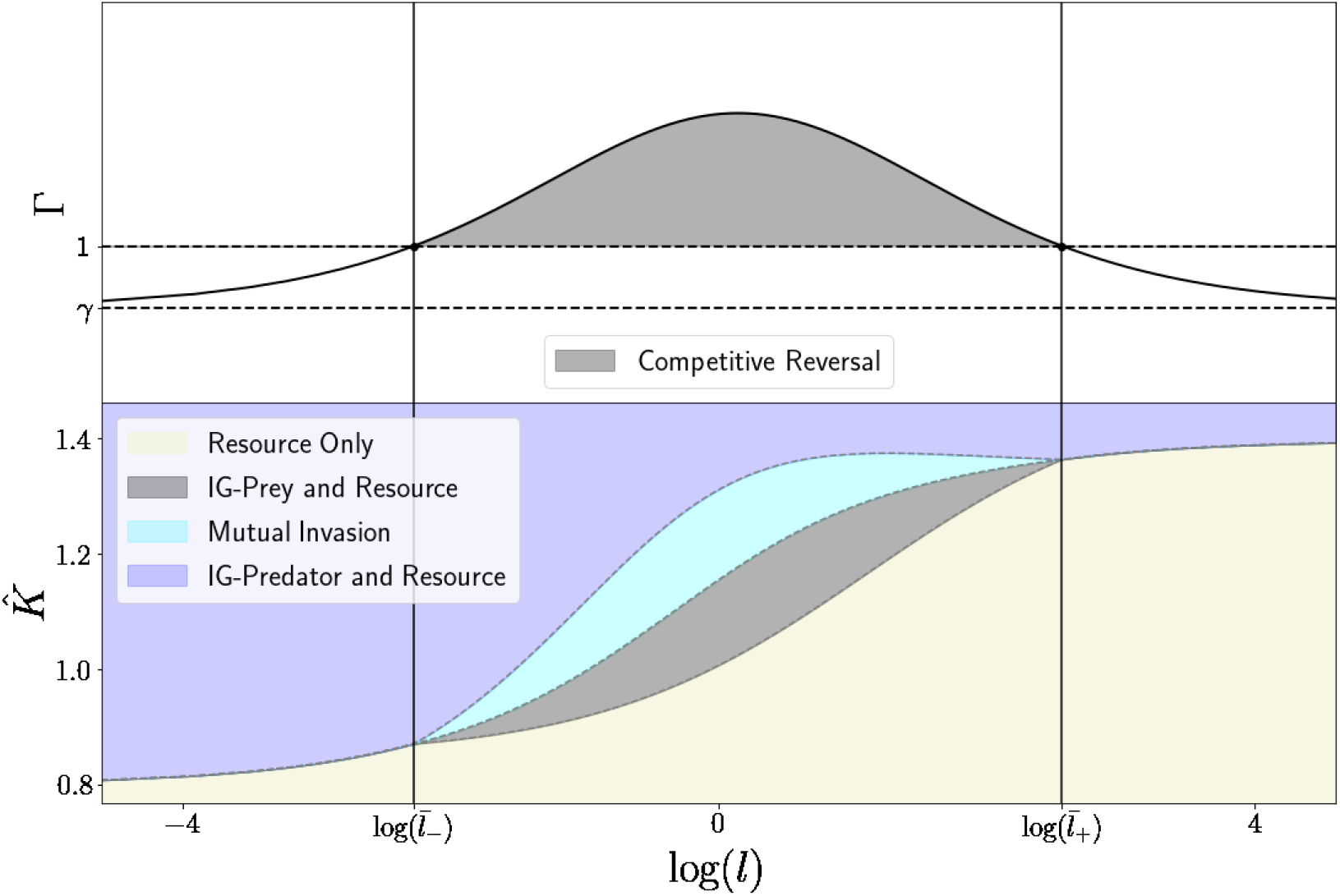
**Top**: Γ as a function of log(*l*). The gray colored region indicates a competitive reversals in favor of IG prey. **Bottom**: Invasibility regimes for different values of 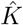. The blue region is bounded bellow by 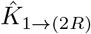 for 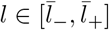, and by 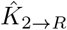 for other values of *l*, cyan region is comprised by 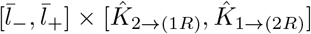, gray region is comprised by 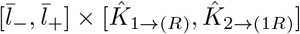, the pastel region is the complementary in the (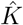, log(*l*)) parameter space. **Parameters used**: *α*_1_ = *α*_2_ = 0.25, *β*_1_ = *β*_2_ = 0.45, *b*_11_ = *b*_12_ = 0.8, *b*_21_ = 0.85*b*_22_ = 0.6, *δ*_11_ = *δ*_12_ = 0.7 *δ*_21_ = 0.85*δ*_22_ = 0.5, *r*_1_ = *r*_2_ = 1, 2*D*_1_ = *D*_2_ = 2*D*_*R*_ = 1.4, *α*_1_ = *α*_*R*_ = 1 − *α*_2_ = 0.45

To compare the approximation in 9 with numerical results, we plot the maximum population densities in figure 10. For that, we let 10 pairs of patches 1 and 2 of equal size (*l*_1_ = *l*_2_ = 1) in order to investigate whether mutual invasibility regimes were indeed observed, and vary the carrying capacities in each patch equally. The thresholds found via homogenization technique are quite close to the ones observed numerically, and mutual invasibility showed to lead to coexistence between IG prey and IG predator.

**Fig. 10:**
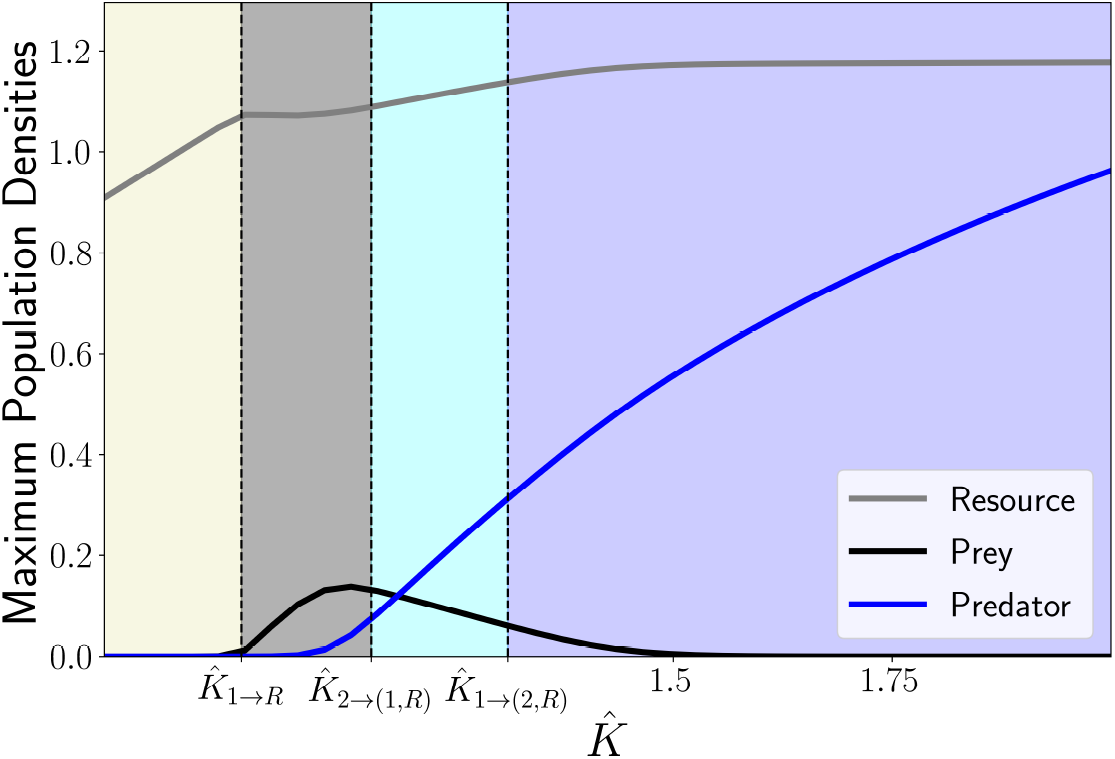
Numerically obtained maximum population densities for *l* = 1. The colored regions and parameters used are the same as in figure 9.

Now, in the case 1 *< γ < w*(*q*_*R*_, *l*\*, *η*_1_, *η*_2_) *>* and (*q*_*r*_, *δη*) ∈ ℋ_*h*_, we have figure 11. The regions of mutual invasibility (cyan colored) and IG prey dominance (gray colored) only exist outside the range 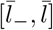, where Γ *>* 1 in the top plot. Inside the interval 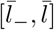, only IG predator is able to invade granted resource carrying capacity is high enough (blue colored), where Γ *<* 1, while for low values of carrying capacity only resource is established in the landscape (pastel colored). This depicts a competitive reversal in favor of IG predator.

**Fig. 11:**
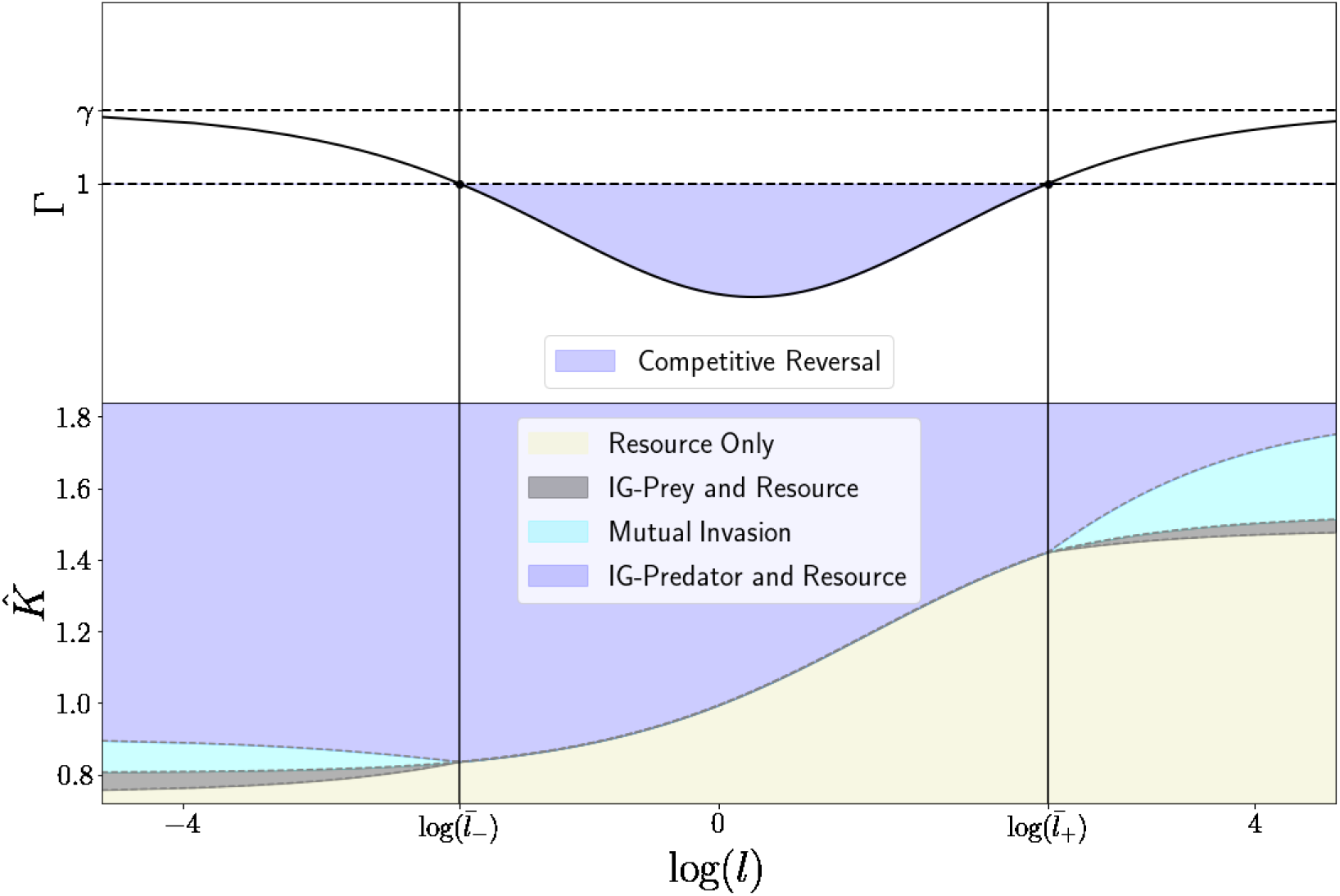
**Top**: Γ as a function of log(*l*). The gray colored region indicates a competitive reversals in favor of IG prey. **Bottom**: Invasibility regimes for different values of 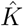. The blue region is bounded bellow by 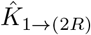 for 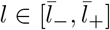, and by 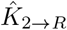 for other values of *l*, cyan region is comprised by 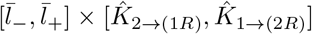, gray region is comprised by 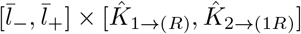, the pastel region is the complementary in the (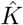, log(*l*)) parameter space. **Parameters used**: *γ* = 1.05, *α*_1_ = *α*_2_ = 0.25, *β*_1_ = *β*_2_ = 0.45, *b*_11_ = *b*_12_ = 0.8, *b*_21_ = 0.85*b*_22_ = 0.6, *δ*_11_ = *δ*_12_ = 0.7 *δ*_21_ = 0.85*δ*_22_ = 0.5, *r*_1_ = *r*_2_ = 1, 2*D*_1_ = *D*_2_ = 2*D*_*R*_ = 1.4, *α*_1_ = *α*_*R*_ = 1 − *α*_2_ = 0.45

Again, we compare the approximation in 11 with numerical results in figure 12. For that, we let 10 pairs of patches 1 and 2 of equal size (*l*_1_ = *l*_2_ = 1) in order to investigate whether exclusion regimes were indeed observed, and vary the carrying capacities in each patch equally, as before. The IG predator threshold found via homogenization technique is quite close to the one observed numerically.

**Fig. 12:**
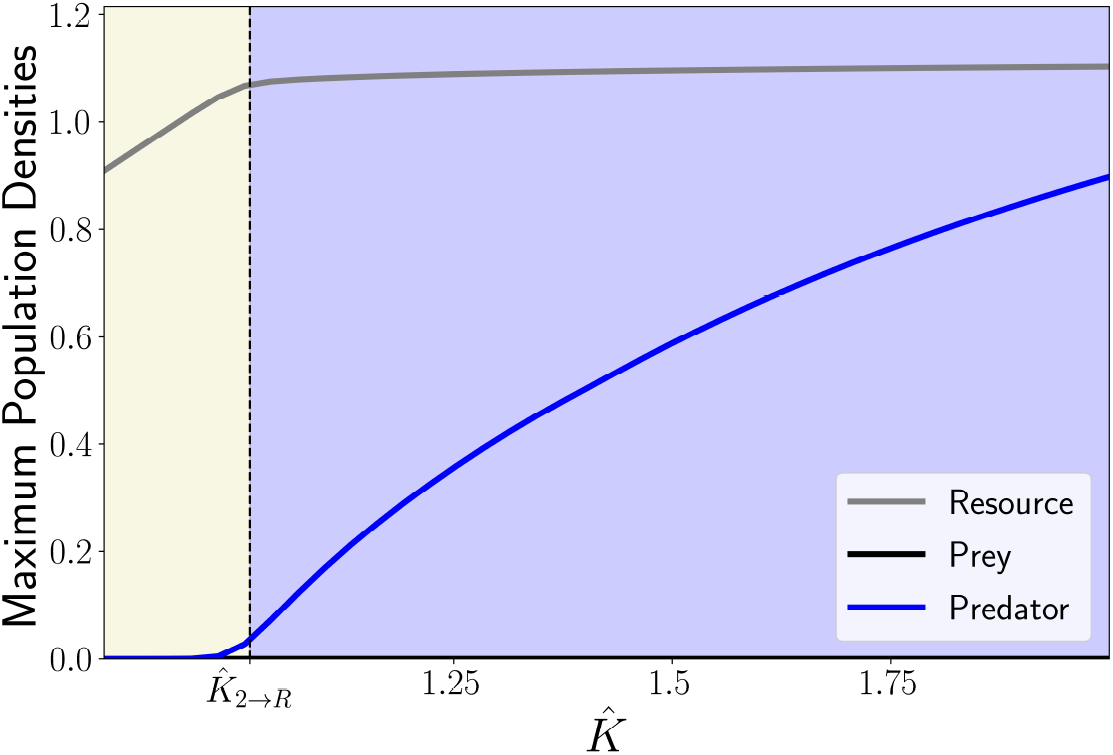
Numerically obtained maximum population densities for *l* = 1. The colored regions and parameters used are the same as in figure 11.

## 4 Discussion

In this work we analyzed of Intraguild Predation communities in homogeneous and heterogeneous landscapes, studying consumer species invasion dynamics. In an homogenous landscape, we recover invasibility conditions as expected in Holt and Polis (1997), while also numerically verified that speeds of invasion are linearly determinate. In heterogeneous environments, using an approximation technique, we found competitive reversals between IG prey and IG predators modulated by multiple factors.

In an homogeneous landscape, we have four possible invasion regimes for when the IG prey is the best exploitative competitor. First, neither of the consumers are able to invade, given a really low carrying capacity, then, for higher carrying capacities we have three distinct ranges: IG prey is able to invade, mutual invasibility between IG prey and IG predator, and IG predator invasions leading to IG prey exclusion and preventing IG prey invasion. For the case where IG predator is the best consumer, we only find two regimes, either no consumer invades for really low carrying capacities, or IG predator invades for large enough carrying capacities, excluding IG prey whenever it is also present in the landscape, or also preventing IG prey invasion. This is the classical result of Holt and Polis (1997), now revisited in the form of invisibility analysis of spatially structured populations.

Our numerical analysis of the homogeneous landscape model suggests that the different speeds of invasion are linearly determined for a large range of parameter values. This was somehow expected, because invasions in consumer-resource models show linearly determinated speed as well (Petrovskii and Malchow, 2000, 1999; Lewis et al., 2016). We show that whenever a successful invasion leads to a shift from the resident community fixed point to a different stable fixed point (in the not spatially structured model sense), we usually have traveling wave solutions connecting these fixed points. We also show that dynamical stability regions can be formed when the coexistence fixed point is unstable, but leave the precise conditions in which to such dynamical stability occur for future research, possibly using the same analysis as in (Malchow and Petrovskii, 2002), which gets slightly more complicated in a three species system.

We have chosen linear functional responses for predator-prey dynamics, for both consumer-consumer or consumer-resource predation. However, if consumer-resource relations are type II Holing functions (Holling, 1959), the single consumer and resource equilibria can be unstable and present oscillations, allowing for coexistence among two consumers without intraguild predation relations (Klausmeier and Tilman, 2002; Armstrong and McGehee, 1980). The invasibility analysis in this scenario, however, gets much more complicated, and studying invasibility criteria, invasion speeds and regime shifts both with and without IGP relations is a possible venue for future research.

The same invasibility regions found in homogeneous landscapes are found in periodic landscapes as well. However, they depend on multiple factors, and competitive reversals might occur. Competitive reversals have also been observed in models for interference competition in periodic landscapes in Maciel et al. (2018), and depend exclusively on patchiness, *l*, and competitor species movement behaviors. Here we show that, in exploitative competition, resource patch preference is a key factor in order for competitive reversals to occur, either favoring IG prey or IG predator, whichever has its patch preference more aligned with resource’s or has a lower mortality rate in resource species favored patch, thereby facilitating or hindering mutual invasibility regimes. Competitive reversals can also occur if resource is unevenly consumed between patches. When resource is much less consumed where one of the consumers has a lower mortality rate, that consumer is benefited and can possibly overcome the fact of being the worst competitor. Similarly, competitive reversals occur if either of the consumers has a higher patch preference for the patch where resource is less consumed.

The observed competitive reversals show a mechanism of bottom-up regulation of Intraguild Predation communities, based on movement behavior of resource population (Holt and Bonsall, 2017). This allows us to question if top-down regulations, based on predator patch preference, are possible in apparent competition interactions. Consider an invasive generalist predator that induces apparent competition between species of the resident community. In homogeneous landscapes, we expect the classical results of Holt (1977), where prey species coexist at lower densities or that one excludes the other. In heterogeneous landscapes, however, consumer patch preference may shift expected exclusion regimes into coexistence ones and vice-versa, while also shifting exclusion of one prey species to the other. This can be possibly verified in a similar framework as displayed here, following Ovaskainen and Cornell (2003); Maciel and Lutscher (2013) to describe species behavior at patch interface and Yurk and Cobbold (2018); Cobbold et al. (2022) to obtain approximate results, highlighting possible future venues where this framework can be applied.

Our work shows that a landscape composed by different patch types with similar lengths/areas, where species can interact and live, can be either detrimental or beneficial for biodiversity in intraguild predation communities, and has several implications in the context of biological invasions or reintroduction. By mutual invasion facilitation, we have possible coexistence regimes which would be otherwise unattainable. For hindered mutual invasion conditions, expected coexistence regimes might collapse, and IG predator may become dominant even if IG prey is the best competitor and carrying capacities are adequate in each of the patch types isolated.

## 5 Acknowledgments

The authors would like to thank the funding institutions that supported their work. SP is thankful for FAPESP (18/24037-4), RAK for FAPESP (2021/14335-0) and CNPq (315641/2021-5).

## Footnotes

1 There are formal results for competition models, based on cooperative systems, see (Weinberger et al., 2002)

2 when the weaker competitor, by exhibiting a more efficient dispersal behavior than its competitor, can potentially exclude or coexist with it (Cantrell et al., 1998)

3 in fact, *b*_*i*_*/a*_*i*_ is the actual conversion rate, but we address *b*_*i*_ as it for short

4 Here, the choice of logistic growth function reflects a living resource species that actively moves and selects between different patches

## Notes

### Competing Interest Statement

The authors have declared no competing interest.

